# Evidence of hippocampal learning in human infants

**DOI:** 10.1101/2020.10.07.329862

**Authors:** C. T. Ellis, L. J. Skalaban, T. S. Yates, V. R. Bejjanki, N. I. Córdova, N. B. Turk-Browne

## Abstract

The hippocampus is essential for human memory. Thus, memory deficiencies in infants are often attributed to hippocampal immaturity. However, the functionality of the infant hippocampus has never been tested directly. Here we report that the human hippocampus is indeed active in infancy. We recorded hippocampal activity using fMRI while awake infants aged 3-24 months viewed sequences of objects. Greater activity was observed when the order of the sequence contained regularities that could be learned compared to when the order was random. The involvement of the hippocampus in such statistical learning, with additional recruitment of the medial prefrontal cortex, is consistent with findings from adults. These results suggest that the hippocampus supports the important ability of infants to extract the structure of their environment through experience.

---

Memory is at the root of human identity, bridging the present into the past and future, fundamental to personality, relationships, expertise, navigation, and imagination. This ability to store and recall life events (episodic memory) requires a brain region known as the hippocampus (Corkin, 2013). The fact that episodic memory is minimal in infants (Richmond and Nelson, 2009), only becoming detailed and stable later in childhood (Keresztes et al., 2018), and that adults remember very little from infancy (infantile amnesia (Akhtar et al., 2018)), has raised the possibility that the hippocampus may not be functional in human infants (Gómez and Edgin, 2016; Nelson, 1995; Schacter and Moscovitch, 1984). However, this has never been evaluated directly. The hippocampus undergoes structural changes well into adolescence (Arnold and Trojanowski, 1996; Gogtay et al., 2006; Schlichting et al., 2017; Uematsu et al., 2012), but what is its function in infancy?

We test the hypothesis that the human infant hippocampus supports the ability to extract regularities across experiences (Fiser and Aslin, 2002; Saffran et al., 1996). Such statistical learning is critical to cognitive development, for acquiring language (Romberg and Saffran, 2010; Werker et al., 2012) and understanding objects (Smith et al., 2018). This hypothesis is based on evidence from human adults that the hippocampus supports statistical learning in addition to episodic memory (Covington et al., 2018; Schapiro et al., 2014, 2012; Turk-Browne et al., 2009). These two functions are thought to rely on separate hippocampal pathways (Schapiro et al., 2017): The trisynaptic or perforant pathway, represented more in the posterior hippocampus, connects entorhinal cortex to dentate gyrus, CA3, and CA1 to enable pattern separation and rapid episodic encoding. The monosynaptic or temporoammonic pathway, represented more in the anterior hippocampus, connects the entorhinal cortex to CA1 directly and supports the integration of inputs to extract regularities. Anatomical connections in the monosynaptic pathway develop earlier than in the trisynaptic pathway (Hevner and Kinney, 1996; Lavenex and Lavenex, 2013), further supporting the hypothesis that the infant hippocampus is involved in statistical learning.

This hypothesis can only be tested in healthy human infants with functional magnetic resonance imaging (fMRI), because of its unique ability to resolve deep-brain structures like the hippocampus (Ellis and Turk-Browne, 2018). This is a challenging technique to use with awake infants during cognitive tasks, including because of head and body motion, an inability to understand or follow task instructions, and general fussiness. This challenge is evident in the extremely small number of published studies of this type (Biagi et al., 2015; Deen et al., 2017; Dehaene-Lambertz et al., 2002). Here we exploit recently developed methods for awake infant fMRI (Ellis et al., 2020) to provide the first evidence of hippocampal function in human infants. Namely, we show that the infant hippocampus is activated by a learning task that requires encoding and integrating visual experiences.

## Role of infant hippocampus in statistical learning

We collected brain imaging data from 24 sessions with infants aged 3–24 months. We defined anatomical regions of interest (ROIs) using a structural MRI obtained in each session (Fig. 1A). We manually segmented the hippocampus bilaterally from the surrounding medial temporal lobe (MTL) cortex. The volume of the hippocampal ROIs was strongly related to age (left *b*=68.0 mm^3^/month, *r*=0.88, *p*<.001; right *b*=68.5 mm^3^/month, *r*=0.84, *p*<.001), with the hippocampus approximately doubling in volume over this age range (Fig. 1B). Global brain volume increased dramatically with age too (*r*=0.90, *p*<.001), but the change in bilateral hippocampal volume persisted after controlling for this global growth (*r*_partial_=0.44, *p*=.005). This suggests that the hippocampus grows rapidly in size during infancy, at a rate that is faster than average in the brain.

**Fig. 1.**
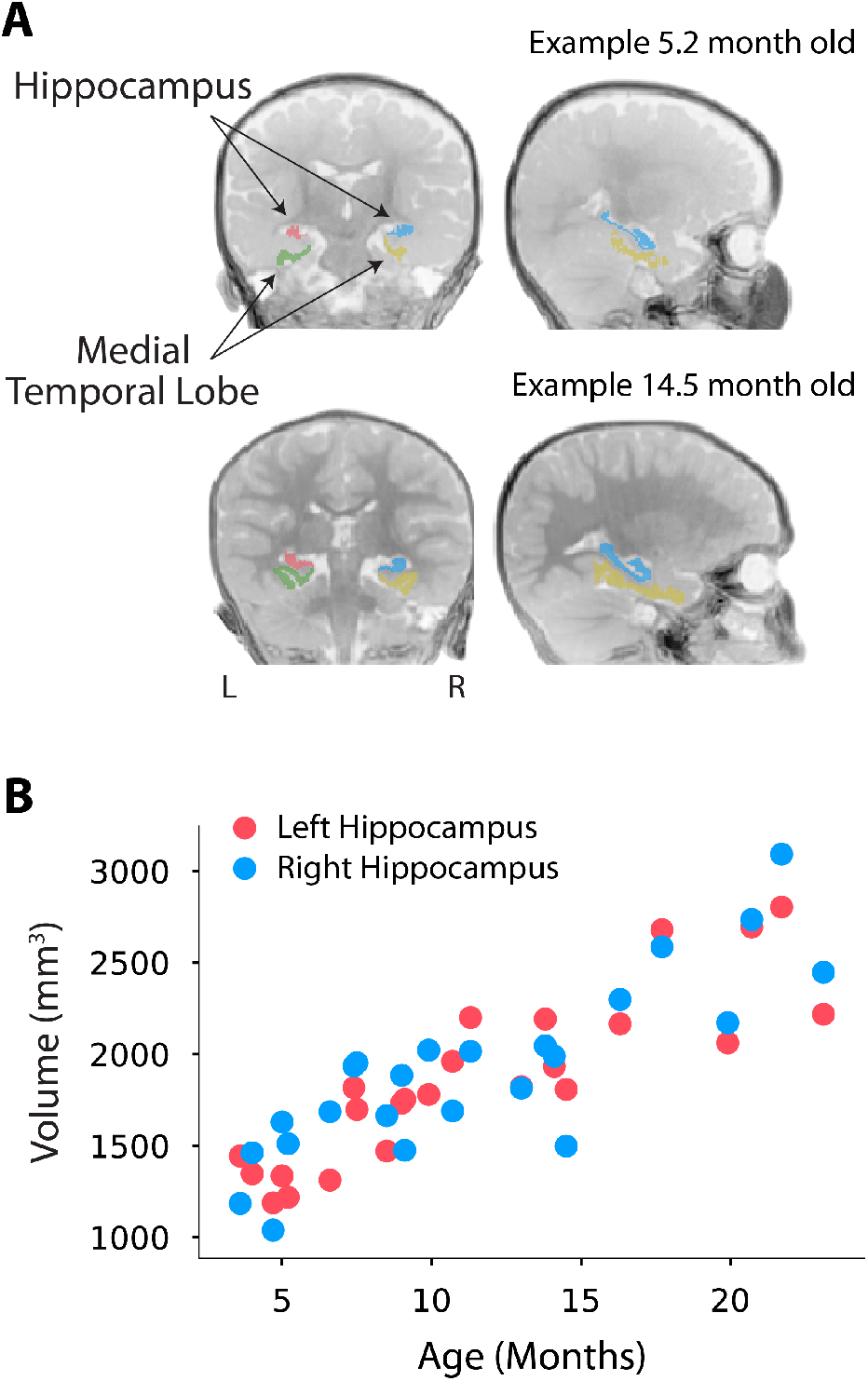
Hippocampal regions of interest. (A) Anatomical segmentation of the infant hippocampus and medial temporal lobe cortex in two representative participants, aged 5.2 months old (top) and 14.5 months old (bottom). (B) Volume of the left and right hippocampus by participant age in months. Each participant is represented by both a red and blue dot at the same age coordinate.

We used fMRI to measure activity in the hippocampus during a statistical learning experiment. Infants viewed continuous sequences of colorful, fractal-like images that appeared dynamically in a looming motion. The sequences were presented in blocks that alternated between Structured and Random conditions (Kirkham et al., 2002). In Structured blocks (Fig. 2A), temporal regularities were embedded in the sequence; fractals appeared in pairs, with the first fractal always followed by the second. In Random blocks (Fig. 2B), there were no regularities in the sequence; rather, all fractals were equally likely to follow each other. Different sets of fractals were used for Structured and Random blocks (counterbalanced across participants), but the fractal set for a given condition was held constant across blocks, as were the pairs generated from the Structured set. Other than the lack of regularities, the Random condition was matched to the Structured condition, including in terms of the number of unique fractals and their frequency across blocks. Any difference in brain activity between Structured and Random blocks can thus be attributed to the presence of regularities in Structured blocks (Turk-Browne et al., 2009). Importantly, representing these regularities required learning: it was necessary to encode and integrate co-occurrences of fractals to extract the pairs from non-paired transitions in the sequence. In other words, because the pairings were arbitrary, at any isolated moment in a Structured block it was impossible to know which fractals were paired; the pairs only exist in the mind of the observer because of the history of how the fractals appeared together earlier in the block or in preceding blocks.

**Fig. 2.**
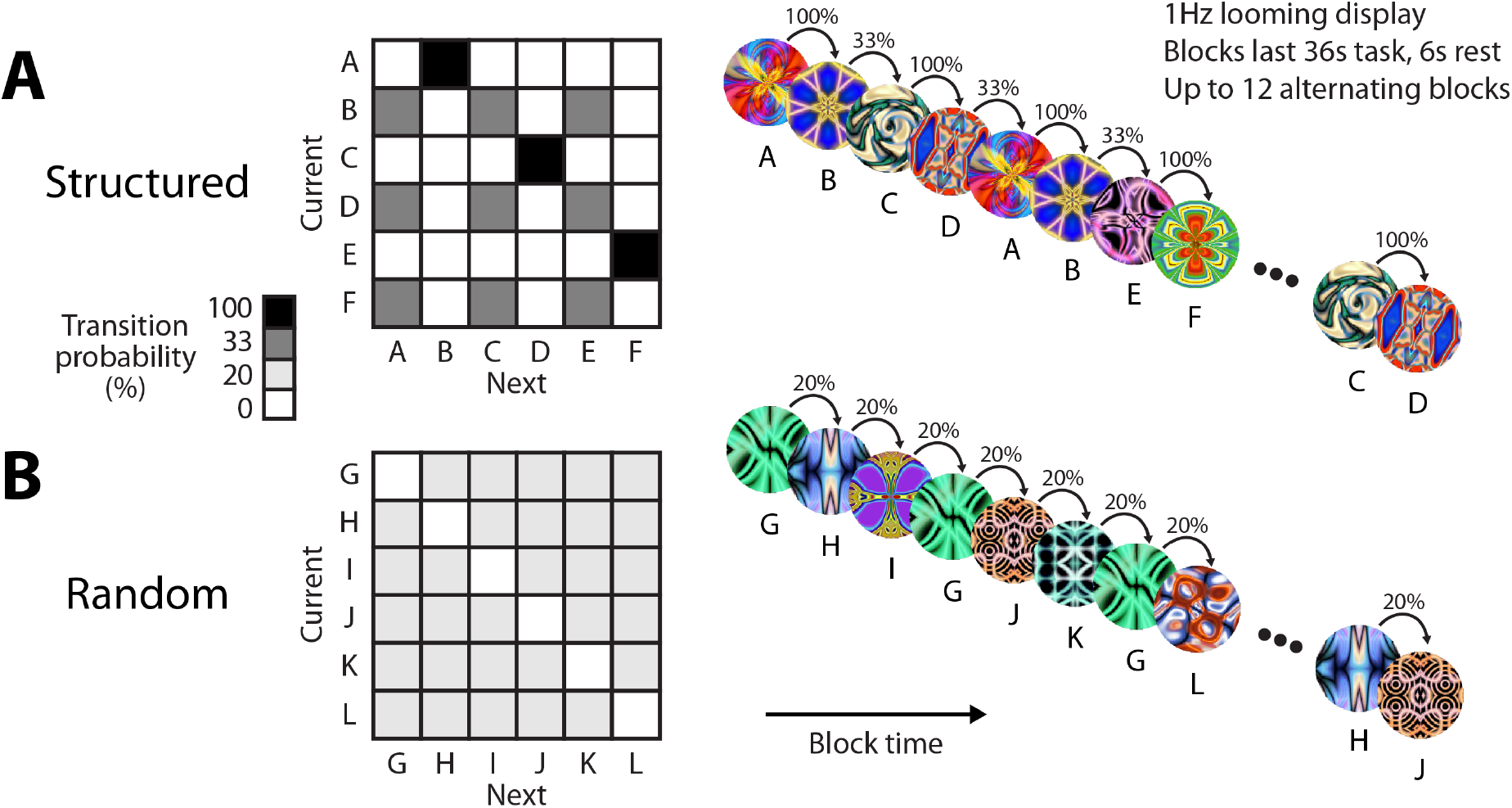
Statistical learning task design. Participants viewed colorful fractals one at a time in blocks. The blocks alternated between a Structured condition and a Random condition within-participant. Different fractals were shown in each condition, but remained consistent over blocks. (A) In Structured blocks, fractals were grouped into three pairs (AB, CD, EF), with the first member of a pair (e.g., A) always followed by the second (e.g., B); this was followed by the beginning of the next pair without interruption. As a result, the pairs could only be learned based on the transition probabilities in the sequence (100% within pair, 33% between pairs). (B) In Random blocks, fractals (G, H, I, J, K, L) appeared in a random order with no back-to-back repetitions. As a result, there was no structure in their transition probabilities (uniform 20%). Because fractals were randomly assigned to the conditions, and individually appeared an equal number of times within and across blocks (to equate familiarity), the conditions differed only in the opportunity for statistical learning. Participants completed up to 12 blocks and usable blocks were split into the first and second half of exposure.

To capture this learning over time, we divided the blocks in each condition into the first half of exposure (when we expected less evidence of learning) and the second half of exposure (when we expected more robust learning effects). We then calculated the difference in blood-oxygenation level dependent (BOLD) response in the bilateral hippocampus between Structured and Random blocks (Fig. 3A). In the first half, there was no difference in hippocampal activity between Structured and Random blocks (M=0.14, CI=[-0.358, 0.656], *p*=.614). However, in the second half, there was significantly greater hippocampal activity in Structured than Random blocks (M=0.67, CI=[0.172, 1.176], *p*=.007). This difference in the second half was larger than in the first half, as revealed by significant interaction between condition and half (M=0.50, CI=[0.028, 0.966], *p*=.037). This learning-related interaction did not differ based on whether infants encountered a Structured or Random block first (M=-0.51, CI=[-1.503, 0.454], *p*=.296), nor did it correlate with the age of the infants (Fig. 3B; *r*=-0.03, *p*=.893). The lack of an age relationship did not reflect a general inability to resolve such relationships in our sample, as the volume of the hippocampus reliably increased over this interval (Fig. 1). These findings suggest that from as young as three months old, the hippocampus is able to support statistical learning. This represents the first evidence of task-related activity in the hippocampus of human infants to our knowledge.

**Fig. 3.**
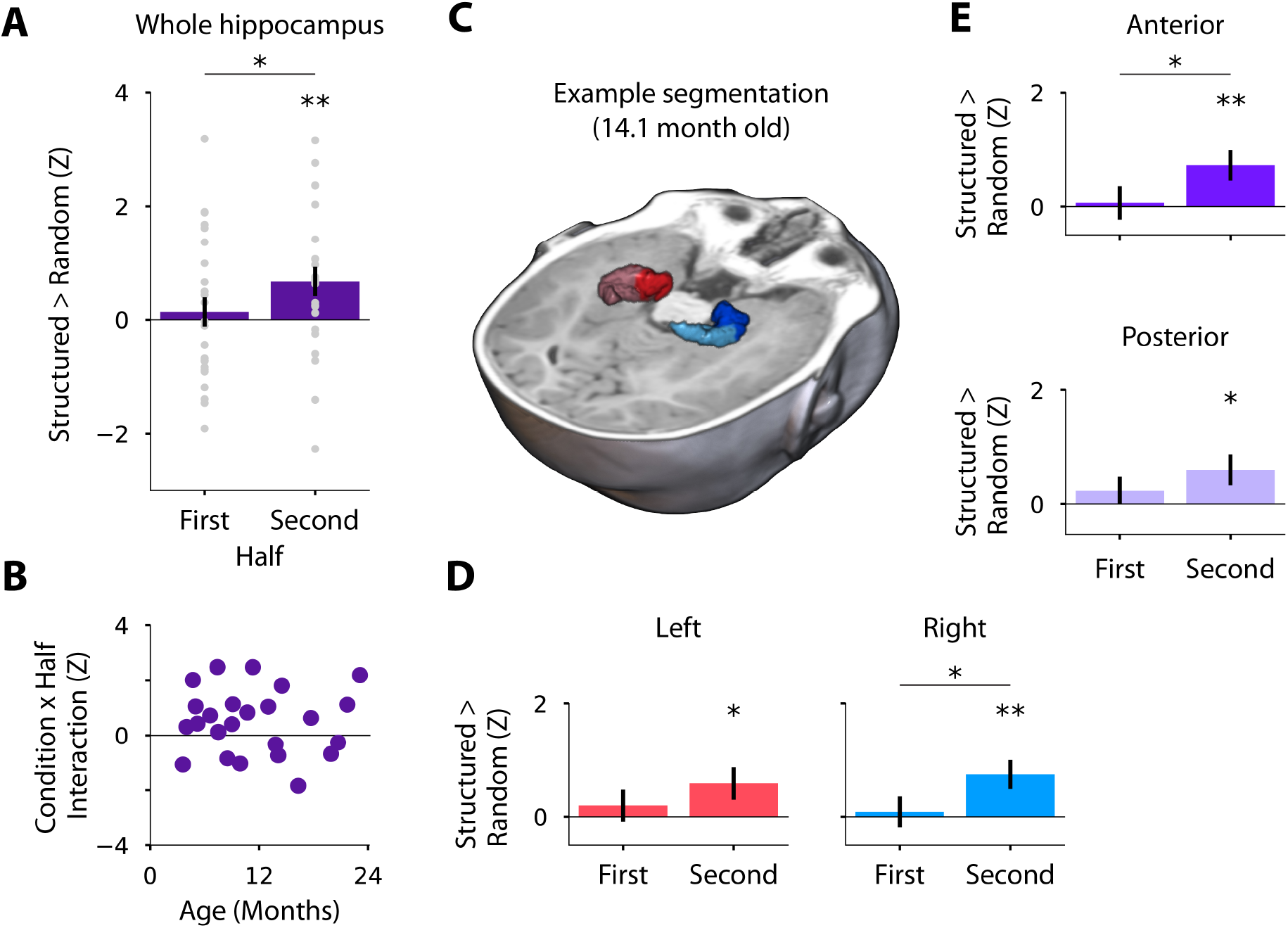
Neural evidence of statistical learning in the infant hippocampus. (A) Mean difference in normalized parameter estimates of BOLD activity between Structured and Random blocks in bilateral hippocampus. A reliable difference emerged by the second half, which was significantly greater than in the first half. Each gray dot is one participant. (B) Using the interaction between condition and half as a metric of hippocampal statistical learning, there was no relationship with participant age in months. (C) Three-dimensional rendering of an example hippocampal segmentation (14.1 month old). Mean difference in BOLD activity between Structured and Random blocks in (D) left (red) and right (blue) hippocampus, and in (E) anterior (dark) and posterior (light) hippocampus. Error bars reflect standard error of the mean across participants within half. * indicates *p*<.05, ** indicates *p*<.01.

## Functional divisions within the hippocampus

We hypothesized that the hippocampus is involved in infant statistical learning partly because of the early development of the monosynaptic pathway from entorhinal cortex to CA1 (Hevner and Kinney, 1996; Lavenex and Lavenex, 2013; Schapiro et al., 2017). There are no established protocols for segmenting hippocampal subfields in infants that would allow us to directly evaluate the role of CA1, and so instead we used the longitudinal axis of the hippocampus as a proxy. Namely, the anterior hippocampus contains more of CA1 than the posterior hippocampus (Malykhin et al., 2010), and so we predicted clearer evidence of statistical learning in the anterior hippocampus (Fig. 3C). Indeed, whereas the anterior hippocampus showed no difference between Structured and Random blocks in the first half (M=0.07, CI=[-0.489, 0.648], *p*=.846), there was a robust difference in the second half (M=0.73, CI=[0.202, 1.257], *p*=.006) and a significant interaction between condition and half (M=0.58, CI=[0.091, 1.063], *p*=.018). The posterior hippocampus again showed no difference in the first half (M=0.23, CI=[-0.216, 0.727], *p*=.328), but the difference in the second half was numerically weaker than in the anterior hippocampus (M=0.60, CI=[0.062, 1.112], *p*=.028) and the interaction did not reach significance (M=0.39, CI=[-0.106, 0.879], *p*=.119); the interaction in posterior was not significantly weaker than in anterior (M=0.19, CI=[-0.086, 0.477], *p*=.174).

In addition to subdividing the longitudinal axis of the hippocampus, we also separated the hippocampus into left and right hemispheres. Adult fMRI studies have reported statistical learning effects more consistently in the right hippocampus (Schapiro et al., 2012; Turk-Browne et al., 2009). This same pattern was found in infants, with numerically stronger evidence of statistical learning in the right hippocampus (first half: M=0.09, CI=[-0.428, 0.635], *p*=.759; second half: M=0.75, CI=[0.243, 1.243], *p*=.003; interaction: M=0.60, CI=[0.116, 1.090], *p*=.013) than in the left hippocampus (first half: M=0.20, CI=[-0.345, 0.754], *p*=.500; second half: M=0.59, CI=[0.021, 1.132], *p*=.043; interaction: M=0.38, CI=[-0.135, 0.890], *p*=.155); the interaction in left was not significantly weaker than in right (M=0.22, CI=[-0.106, 0.574], *p*=.207). Thus, the overall pattern of learning-related signals across longitudinal and hemispheric axes of the infant hippocampus is consistent with primate anatomy (Hevner and Kinney, 1996; Lavenex and Lavenex, 2013), computational models (Schapiro et al., 2017), and adult function (Schapiro et al., 2012; Turk-Browne et al., 2009).

## Timecourse of hippocampal involvement

Splitting the fMRI data into the first and second half of exposure was an attempt to capture learning over time while retaining enough blocks per time bin to estimate stable effects. We also examined learning over time more continuously at the block level (Fig. S1). Adopting a supersubject approach, we pooled usable blocks across participants and assessed statistical significance with bootstrap resampling. The difference between Structured and Random blocks was largest and only statistically significant in the fifth and sixth blocks (of six). In other words, evidence of statistical learning emerged after approximately two minutes of exposure to Structured blocks (four blocks of 36 s).

The amount of exposure needed to obtain neural evidence of statistical learning is consistent with the duration of classic behavioral studies of infant statistical learning (Kirkham et al., 2002; Saffran et al., 1996). This suggests that fMRI can serve as a sensitive, converging measure of infant cognition, even for relatively short task designs. An important limitation of the current study is that we did not obtain a behavioral measure of statistical learning that could be directly related to the fMRI findings. Nevertheless, the design of our study, in which Random blocks carefully controlled for all aspects of Structured blocks other than the presence of regularities to be learned, allows us to attribute the observed neural differences to statistical learning.

## Engagement of neocortical systems

The focus of this study was on examining the function of the infant hippocampus, with the hypothesis that it supports statistical learning. However, we also compared Structured and Random blocks in the surrounding MTL cortex and found weak evidence of statistical learning (Fig. S2). Given that MTL cortex is anatomically adjacent and a larger ROI, this highlights the specificity of our findings in the hippocampus. We additionally performed exploratory voxelwise analyses across the whole brain with data aligned to standard space across participants (Fig. S3). The key learning interaction observed in the hippocampus between condition (Structured vs. Random) and half (second vs. first) was found only in medial prefrontal cortex (mPFC; corrected *p*=.048, 116 voxels, MNI: -5, 53, 3).

This involvement of mPFC in infants is striking given the dramatic changes in frontal lobe anatomy over development (Matsuzawa et al., 2001). In adults, mPFC strongly interacts with the hippocampus during memory formation, facilitating encoding based on related past experiences (i.e., schemas) to promote memory integration (Schlichting et al., 2015) and consolidation (Tse et al., 2011). Indeed, mPFC has been linked to gradual statistical learning over days and weeks in rodents (Richards et al., 2014). It remains to be seen whether this mechanism contributes to rapid statistical learning over minutes in human infants, as tested here. An important limitation of the current study is the inability of fMRI to distinguish whether evidence of statistical learning in the hippocampus originates in the hippocampus or is a reflection of processing in the mPFC, given their connectivity.

## Open questions and theoretical implications

The key finding of this study is that activity in the hippocampus of human infants increases through exposure to regularities. This activity may correspond to different stages of statistical learning. It could reflect the process of extracting regularities during learning, with differences emerging in the second half because a certain amount of exposure was needed to compute the transition probabilities between fractals and represent the pairs. Alternatively, the hippocampal activity could reflect the impact of known regularities on other processes after learning is complete, including on perception of the fractals, segmentation of the sequence, recognition of the pairs, and/or prediction based on transition probabilities. In addition to clarifying at which of these stages the hippocampus participates in infant statistical learning, future research will be needed to determine whether this role is necessary for behavioral expression of statistical learning. This will be difficult to test in infants, but studies in adult patients with hippocampal damage suggest that the hippocampus may in fact be necessary for normal statistical learning behavior (Covington et al., 2018; Schapiro et al., 2014).

The involvement of the infant hippocampus in statistical learning has implications for theories of memory. For example, according to complementary learning systems (McClelland et al., 1995), episodic memory is a precursor to statistical learning. The hippocampus rapidly encodes individual experiences and then, through a process of consolidation, the neocortex gradually generalizes across these episodic memories to extract regularities. Infants present a conundrum for this framework: they show robust statistical learning (Kirkham et al., 2002; Saffran et al., 1996) despite impoverished episodic memory (Akhtar et al., 2018; Keresztes et al., 2018; Richmond and Nelson, 2009). A recent update to complementary learning systems (Schapiro et al., 2017) provides a potential resolution, at least for the rapid form of statistical learning in our study. Neural network simulations showed that such statistical learning can occur within the hippocampus itself in a way that bypasses the circuitry for episodic memory. Thus, the hippocampus may support statistical learning in infants, as reported in this study, before it can support episodic memory. It also remains possible that episodic memory is more developed in infants than currently thought—consistent with recent rodent work (Farooq and Dragoi, 2019; Guskjolen et al., 2018)—such that the hippocampal statistical learning we report may in fact be dependent upon episodic memory. Future research could address these possibilities by using fMRI with awake infants to capture sensitive neural measures of episodic memory functions in the hippocampus, including pattern separation, relational binding, and pattern completion.

To conclude, we present the first evidence that the hippocampus is recruited for learning in human infants. This demonstrates that brain systems used for learning throughout the lifespan can be available from some of the earliest stages of life. In turn, this provides a starting point for understanding how the human brain supports the prodigious amount of learning that occurs during infancy, establishing building blocks critical for subsequent growth and education.

## Acknowledgments

We thank all of the families who participated; K. Armstrong, C. Greenberg, J. Bu, L. Rait, and the entire Yale Baby School team for recruitment, scheduling, and administration; H. Faulkner, Y. Braverman, J. Fel, and J. Wu for help with gaze coding; B. Sherman for advice on hippocampal segmentation; and N. Wilson, R. Lee, L. Nystrom, N. DePinto, and R. Watts for technical support. We are grateful for internal funding from the Department of Psychology and Princeton Neuroscience Institute at Princeton University and from the Department of Psychology and Faculty of Arts and Sciences at Yale University. N.B.T-B. was further supported by the Canadian Institute for Advanced Research.

## Author contributions

C.T.E., N.I.C., V.R.B., & N.B.T-B. initially created the protocol. All authors collected the data. C.T.E., L.J.S., T.S.Y., & N.B.T-B. developed the pipeline. C.T.E., L.J.S., & T.S.Y. performed the analyses. C.T.E., & N.B.T-B. wrote the initial draft of the manuscript. All authors contributed to the editing of the manuscript.

## Competing interests

Authors declare no competing interests.

## Correspondence

Please address correspondence and requests for materials to.

## Methods

### Participants

Data from 24 sessions with infants aged 3.6 to 23.1 months (M=11.6, SD=5.8; 14 female) met our minimum criteria for inclusion of six usable task blocks with at least one pair of Structured and Random blocks in each of the first and second halves of exposure (M=11.8 total blocks, M=9.8 usable blocks). This sample does not include data from 11 sessions with enough blocks only prior to exclusions for head motion, eye gaze, and counterbalancing (M=9.8 total blocks, M=3.5 usable blocks), or from 44 sessions without enough blocks even prior to exclusions (M=3.6 total blocks) where the infant instead participated in other experiments. In the final sample, five infants provided two sessions of usable data and one infant provided three. These sessions occurred at least one month apart (range=1.1–9.3) and so the data were treated separately, similar to prior work (Deen et al., 2017). Of the 24 sessions, six were collected at the Scully Center for the Neuroscience of Mind and Behavior at Princeton University, four were collected at the Magnetic Resonance Research Center (MRRC) at Yale University, and 14 were collected at the Brain Imaging Center (BIC) at Yale University. Refer to Table S1 for information on each participant. Parents provided informed consent on behalf of their child. The study was approved by the Institutional Review Board at Princeton University and the Human Investigation Committee at Yale University.

### Data acquisition

Data were acquired with a Siemens Skyra (3T) MRI at Princeton University and a Siemens Prisma (3T) MRI at both sites at Yale University, in all cases with the 20-channel Siemens head coil. Anatomical images were acquired with a T1-weighted PETRA sequence (TR_1_=3.32ms, TR_2_=2250ms, TE=0.07ms, flip angle=6°, matrix=320×320, slices=320, resolution=0.94mm iso, radial slices=30000). Functional images were acquired with a whole-brain T2* gradient-echo EPI sequence (Princeton and Yale MRRC: TR=2s, TE=28ms, flip angle=71°, matrix=64×64, slices=36, resolution=3mm iso, interleaved slice acquisition; Yale BIC: identical except TE=30ms, slices=34).

### Procedure

Conducting fMRI research with awake infants presents many challenges. We have described and validated our protocol in detail in a separate methods paper (Ellis et al., 2020). In brief, families visited the lab prior to their first scanning session for an orientation session. This served to acclimate the infant and parent to the scanning environment. Scanning sessions were scheduled for a time when the parents felt the infant would be calm and happy. The infant and parent were extensively screened for metal. Hearing protection was applied to the infant in three layers: silicon inner ear putty, over-ear adhesive covers, and ear muffs. The infant was placed on the scanner bed, on top of a vacuum pillow that comfortably reduced movement. The top of the head coil was not used because the bottom elements provided sufficient coverage of the smaller infant head. This created better visibility for monitoring infant comfort and allowed us to project stimuli onto the ceiling of the bore directly above the infant’s face using a custom mirror system. A video camera (Princeton and Yale MRRC: MRC 12M-i camera; Yale BIC: MRC high-resolution camera) recorded the infant’s face during scanning for monitoring and eye tracking.

When the infant was calm and focused, stimuli were shown in Matlab using Psychtoolbox (http://psychtoolbox.org). The stimuli were colorful, fractal-like images used previously in studies of statistical learning in adults (Hindy et al., 2016; Schapiro et al., 2012). Images appeared every 1s, looming in size from 2.4° at onset to 14.6° degrees at offset (Kirkham et al., 2002). Each block contained 36 images presented sequentially one at a time in a unique order, followed by 6s of rest with the screen blank.

Blocks alternated between Structured and Random conditions (Fig. 2). Which condition appeared first was assigned randomly. In the Structured condition, six fractals (A-F) were organized into three pairs (AB, CD, EF). The sequence of each block was generated by randomly inserting six repetitions of each pair. The first member of a pair (A, C, E) was always followed by the second (B, D, F, respectively) resulting in a transition probability of 1.0. After the second member of a pair, another pair appeared, resulting in a transition probability of 0.33 on average. In the Random condition, six different fractals (G-L) were presented individually. The sequence of each block was generated by randomly inserting six repetitions of each fractal, avoiding back-to-back repetitions of the same fractal. This resulted in a uniform transition probability of 0.20 on average. The six fractals in each condition were consistent across all blocks of that condition. For participants who attempted the experiment in more than one session, different stimuli were used across sessions.

### Gaze coding

Infant gaze was coded offline by two or more coders (M=2.65) blind to the block condition. The coders determined whether the gaze was on-screen, off-screen (i.e., blinking or looking away), or undetected (i.e., out of the camera’s field of view or obscured by a hand or other object). Across coders, every video frame was coded at least once. The frame rate and resolution varied by camera and site, but the minimum rate was 16Hz and we always had sufficient resolution to identify the eye. The coded category for each frame was determined as the mode of a moving window of five frames centered on that frame across all coder reports. In case of a tie, the modal response from the previous frame was used. The coders were highly reliable: when coding the same frame, coders reported the same response on 93% (SD=6%; range across participants=73–99%) of frames. Infants included in the final sample looked at the stimulus 89% of the time on average (range=80.3–97.3%). Blocks were excluded if the eyes were off-screen for 50+% of the block. One participant did not have eye-tracking data due to a technical problem but real-time monitoring confirmed that their eyes were open and attending to the stimulus for at least 50% of each block.

### Preprocessing

Individual runs were preprocessed using FEAT in FSL (https://fsl.fmrib.ox.ac.uk/fsl), with modifications optimized for infant data. We discarded three volumes from the beginning of each run, in addition to the volumes automatically discarded by the EPI sequence. Blocks were stripped of any excess burn-in or burn-out volumes beyond the 3 TRs (6s) of rest after each block. Pseudo-runs were generated if other experiments, not discussed here, were initiated in a run with the data of interest (sessions with a pseudo-run, N=12). Blocks were sometimes separated by long pauses (>30s) within a session because of a break outside of the scanner, because an anatomical scan was collected, or because of intervening experiments (N=7; M=636.7s break; range=115.4–1545.1s). The reference volume for alignment and motion correction was chosen as the ‘centroid’ volume with the minimal Euclidean distance from all other volumes. The slices in each volume were realigned with slice-time correction. Time-points were excluded if there was greater than 3mm of movement from the previous time-point (M=8.9%, range=0.0–21.3%). We interpolated rather than excised these time-points so that they did not bias the linear detrending (in later analyses these time-points were excised). Blocks were excluded if 50+% of the time-points were excluded. The mask of brain and non-brain voxels was created from the signal-to-fluctuating-noise ratio (SFNR) for each voxel in the centroid volume. The data were spatially smoothed with a Gaussian kernel (5mm FWHM) and linearly detrended in time. The despiking algorithm in AFNI (https://afni.nimh.nih.gov) was used to attenuate aberrant time-points within voxels. For further explanation and justification of this preprocessing procedure, refer to (Ellis et al., 2020).

We registered each run’s centroid volume to the infant’s anatomical scan from the same session. We used FLIRT with a normalized mutual information cost function for initial alignment. Supplemental manual registration was then performed using mrAlign from mrTools (Gardner lab) to fix deficiencies of automatic registration. The preprocessed functional data were aligned into anatomical space but kept in their original spatial resolution (3mm iso). Region of interest (ROI) analyses were performed within this native space of each participant. Whole-brain voxelwise analyses required further alignment of functional data into a standard space. The anatomical scan from each participant was automatically (FLIRT) and manually (Freeview) aligned to an age-specific MNI infant template (Fonov et al., 2011). Combined with alignment of these templates to the adult MNI template (MNI152), the functional data were transformed into standard space. To determine which voxels to consider at the group level, the intersection of brain voxels from all infant participants in standard space was used as a whole-brain mask.

Because runs could contain different numbers of blocks from the Structured and Random conditions, blocks were only retained if they could be paired with a block from the other condition in the same run. This counterbalancing was enforced to ensure an equal amount of data in each condition. The blocks were labeled by the count of how many blocks from that condition had already been seen (henceforth, their ‘seen-count’). For example, if an infant was watching the screen but moving too much in their first Structured block, then remained still in their second Structured block, the first usable block of that condition would be labeled with a seen-count of 2. Blocks were chosen to be paired across conditions so as to minimize the difference in seen-counts (i.e., to match the degree of exposure as best possible).

For an infant to be included, they needed to have at least three blocks from each condition, with at least one block in each condition from blocks 1 to 3 (first half) and at least one block in each condition from blocks 4 to 6 (second half). Using these criteria, the average number of included blocks for the usable participants was 9.8 (SD=1.9, range=6–12), including 5.5 blocks in the first half and 4.3 blocks in the second half on average. There was no correlation between the number of included blocks and age (r=-0.05, *p*=.788). The block order was determined randomly, with 15 participants seeing a Structured block first and 9 participants seeing a Random block first (as reported in the main text, there were no reliable order effects on the neural results).

To account for differences across runs in intensity and variance, the blocks that survived exclusions and balancing across conditions were normalized over time within run using *z*-scoring, prior to the runs being concatenated for further analyses.

### Regions of interest

The main analyses involved manually tracing ROIs in the medial temporal lobe (MTL) based on anatomical landmarks and then assessing evoked BOLD responses across voxels in these anatomical ROIs. To trace the ROIs, we extended a published protocol for MTL segmentation in adults (Aly and Turk-Browne, 2015) with help from protocols for hippocampal segmentation in infants (Gousias et al., 2013). The segmentation demarcated ROIs for the left and right hippocampus, each of which encompassed the subiculum, CA1, CA2/3, and dentate gyrus subfields. We did not individually segment these subfields because of the lack of validated anatomical guidelines for subfield boundaries in infants. For completeness, we also defined ROIs for the left and right MTL cortex, each of which contained the entorhinal, perirhinal, and parahippocampal cortices (again not segmented individually). To examine the reliability of the coder performing the infant segmentations, an expert adult coder segmented two infant participants. Using Dice similarity (Dice, 1945), the consistency of labelling was 0.524 and 0.651 for the two participants across coders, indicating moderate reliability. Fig. 1 shows example ROIs for two infants and the volume of each ROI across participants as a function of age. The anterior hippocampus (volume: M=1973.1 mm^3^, SD=537.5) was defined as the head of the hippocampus, as manually traced (Aly and Turk-Browne, 2015), and the posterior hippocampus (volume: M=1796.4 mm^3^, SD=433.7) was the remainder, including the body and tail. For one participant (4.0 month old), the anatomical scan collected in the same session as the functional data was of insufficient quality for segmentation; we instead used the anatomical scan collected in their next session (at 6.0 months) and aligned the resulting segmentation to their functional data.

### Analysis

For each infant, the volume of left and right hippocampus and MTL cortex ROIs was estimated by counting the number of voxels traced and multiplying by the volume of each voxel (0.82mm^3^). Whole-brain volume was calculated based on the number of voxels in the brain mask generated by applying Freesurfer (https://surfer.nmr.mgh.harvard.edu) to their anatomical scan (Schlichting et al., 2017).

For the main analysis, a general linear model (GLM) was fit to the BOLD activity in each voxel using FEAT in FSL. The GLM contained four regressors: Structured and Random conditions in the first and second half of exposure. Each regressor modeled corresponding task blocks with a boxcar lasting the duration of stimulation convolved with a double-gamma hemodynamic response function. The assignment of blocks to halves was based on the seen-count: blocks with seen-count 1–3 were assigned to the first half and blocks with seen-count 4–6 were assigned to the second half. The six translation and rotation parameters from motion correction were included in the GLM as regressors of no interest. Excluded TRs were scrubbed by including an additional regressor for each to-be-excluded time-point (Siegel et al., 2014). Contrasts of the resulting parameter estimates compared Structured greater than Random conditions separately for the first and second half; an interaction contrast compared the condition differences in the second versus first half. The voxelwise *z*-statistic volumes for these contrasts were extracted for each participant. ROI analyses averaged the *z*-statistics of all included voxels and examined the reliability of these averages at the group level. Whole-brain analyses examined the reliability of the *z*-statistics for each voxel across participants.

Statistical analysis was performed on the ROI data using a non-parametric bootstrap resampling approach (Efron and Tibshirani, 1986). Namely, for each test we sampled 24 participants with replacement 10,000 times, averaging across participants on each iteration to generate a sampling distribution. For null hypothesis testing, we calculated the *p*-value as the proportion of samples whose mean was in the opposite direction from the true effect, doubled to make the test two-tailed. To correct for multiple comparisons in whole-brain analyses, we used threshold free cluster enhancement through the randomise function in FSL, resulting in voxel clusters *p*<.05 corrected. A similar bootstrap resampling procedure was used to statistically evaluate correlations, sampling bivariate data from 24 participants with replacement 10,000 times, and calculating the Pearson correlation (or partial correlation) on each iteration. We calculated the *p*-value as the proportion of samples resulting in a correlation with the opposite sign from the true correlation, doubled to make the test two-tailed.

To perform the timecourse analysis (Fig. S1), we restricted analysis to pairs of Structured and Random blocks with identical seen-counts (as opposed to finding the closest match in the main analysis). This allowed us to separately examine the difference between Structured and Random at each of the 6 ordinal positions. This reduced the number of participants with a sufficient number of usable blocks to 22, and the average number of usable blocks per retained participant to 9.6 (SD=1.9; range=6–12). A GLM was fit to these data with a separate regressor for each block. The parameter estimates were labeled based on each block’s seen-count and contrasted across conditions within the same seen-count. The resulting *z*-statistics were averaged across voxels within each ROI. The same bootstrap resampling approach with 1,000 iterations was used to assess statistical reliability and calculate *p*-values for each ordinal position. An important feature of this approach is that we were able to estimate the timecourse even if individual subjects were missing one or more of the positions. This also takes into account the smaller sample size of participants with later ordinal positions, because the obtained sampling distribution is more variable.

## Data availability

The data, including anonymized anatomical images, manually segmented regions, and both raw and preprocessed functional images will be released on Dryad upon publication.

## Code availability

The code for running the statistical learning task can be found here: https://github.com/ntblab/experiment_menu. The code for the general analysis pipeline can be found here: https://github.com/ntblab/infant_neuropipe. The code for performing the specific analyses reported in this paper can be found here: https://github.com/ntblab/infant_neuropipe/tree/StatLearning.

---

**Fig. S1.**
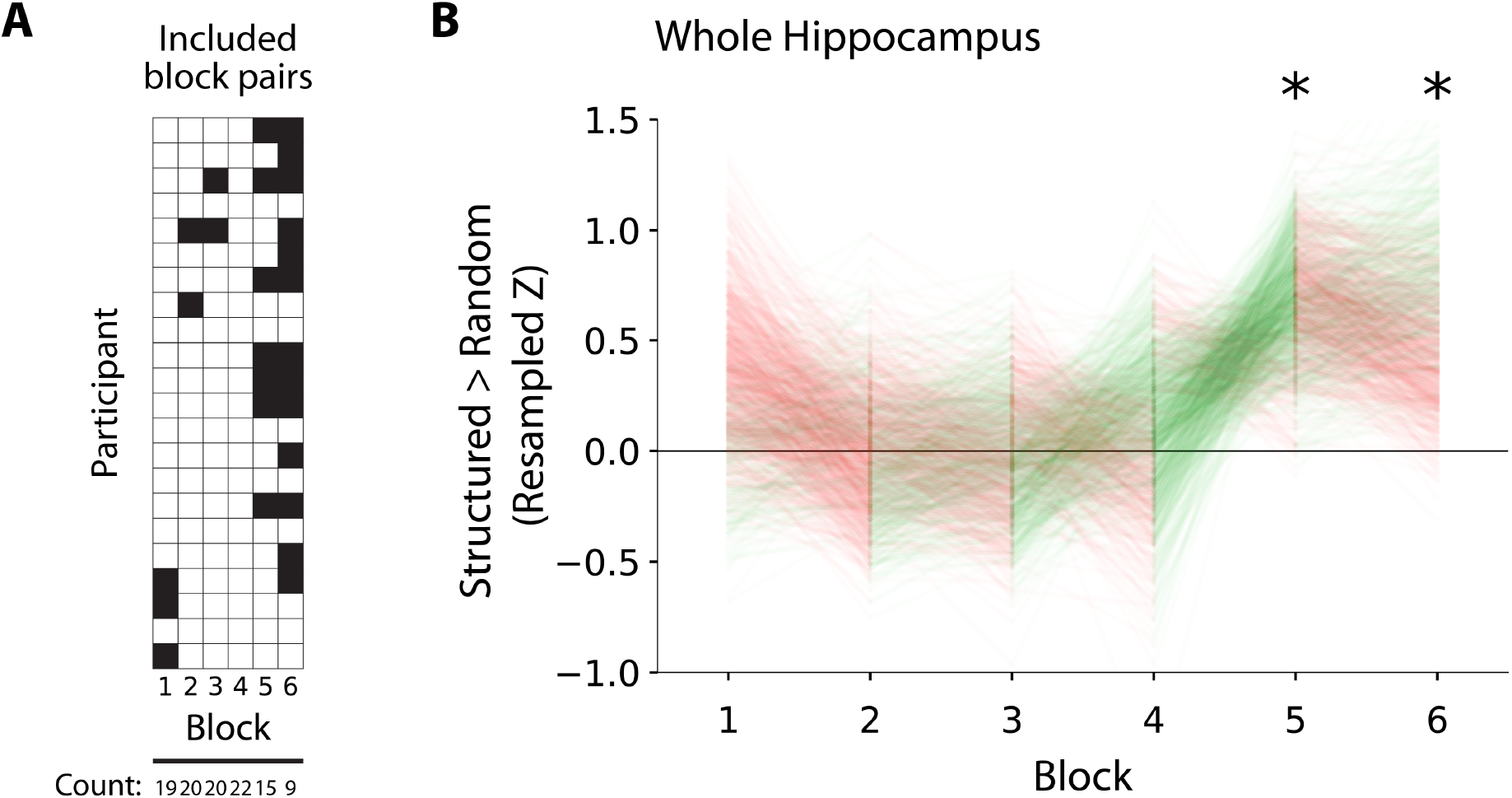
Timecourse of hippocampal statistical learning in infants. (A) To examine learning more continuously, we pooled blocks across participants (rows) based on their ordinal position during exposure (columns). White cells indicate that both the Structured and Random blocks from that ordinal position were usable and included in the timecourse analysis; black cells indicate that the block from one or both conditions are not usable, and therefore neither was included. (B) Difference in BOLD activity between Structured and Random blocks in bilateral hippocampus as a function of ordinal position: block 1, bootstrapped *p*=.362; block 2, *p*=.989; block 3, *p*=.939; block 4, *p*=.661; block 5, *p*=.010; block 6, *p*=.042. Individual lines correspond to bootstrapping iterations and thus convey the sampling distribution. Line color indicates whether the bootstrapped mean increases (green) or decreases (red) from one block to the next on a given iteration. * indicates that the 95% confidence interval does not contain zero.

**Fig. S2.**
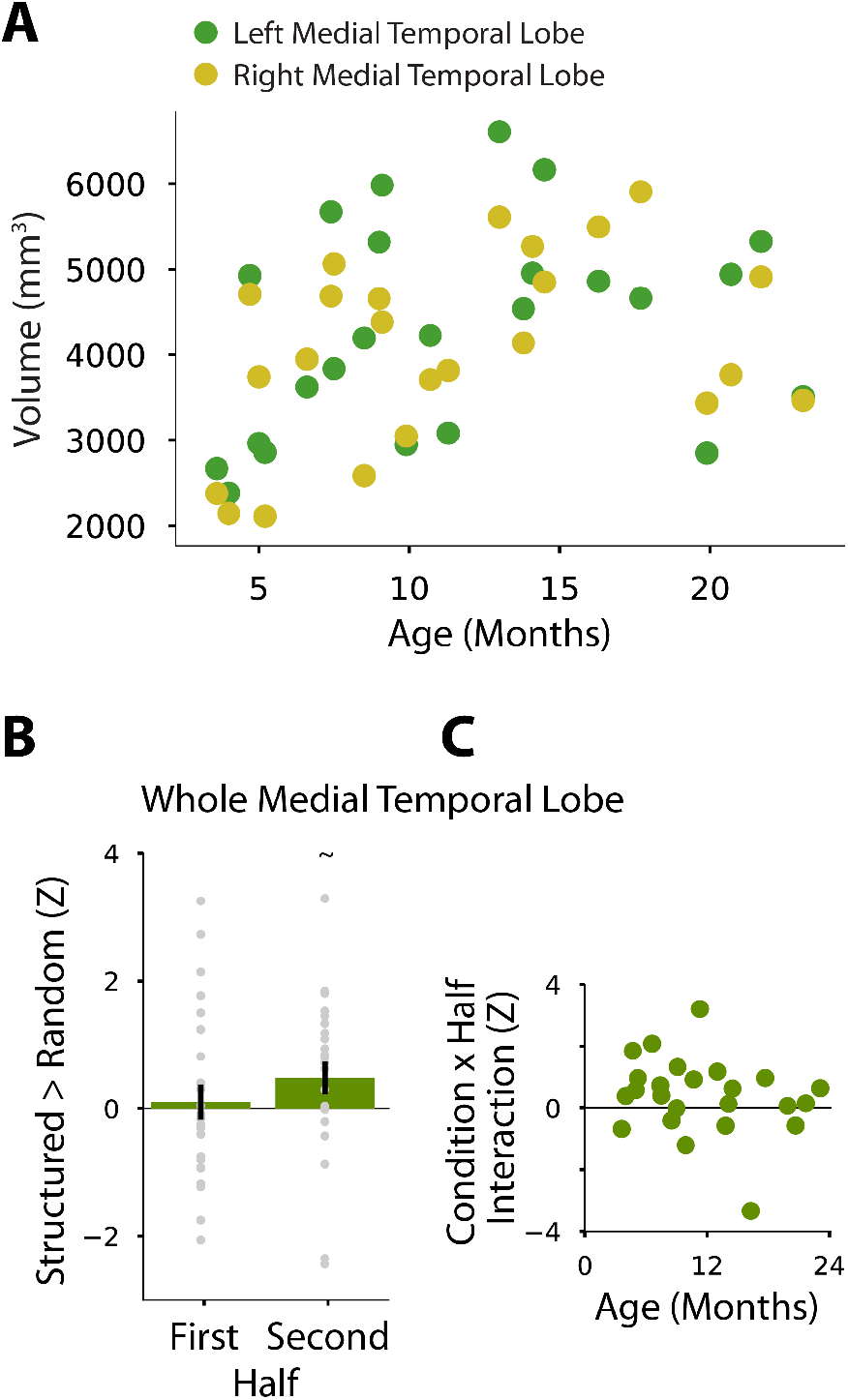
Medial temporal lobe (MTL) cortex. (A) Volume of anatomically segmented left and right MTL cortex by participant age. Unlike the hippocampus (Fig. 1A), there was no reliable relationship between age and volume (left *b*=59.3 mm^3^/month, *r*=0.29, *p*=.142; right *b*=69.2 mm^3^/month, *r*=0.37, *p*=.062). Although the slope values are similar to the hippocampus, the larger size of MTL cortex (M=8371.8 mm^3^, SD=2119.2) compared to the hippocampus (M=3769.4 mm^3^, SD=898.2), means that the proportional growth is much lower. (B) Mean difference in BOLD activity between Structured and Random blocks in bilateral MTL cortex by exposure half. The first half (M=0.10, CI=[-0.423, 0.682], *p*=.726), second half (M=0.48, CI=[-0.023, 0.959], *p*=.059), and interaction between condition and half (M=0.40, CI=[-0.111, 0.875], *p*=.121) did not reach significance. (C) There was no reliable relationship between this interaction and age (*r*=-0.23, *p*=.273). Error bars depict standard error of the mean across participants within half. ∼ indicates *p*<.10.

**Fig. S3.**
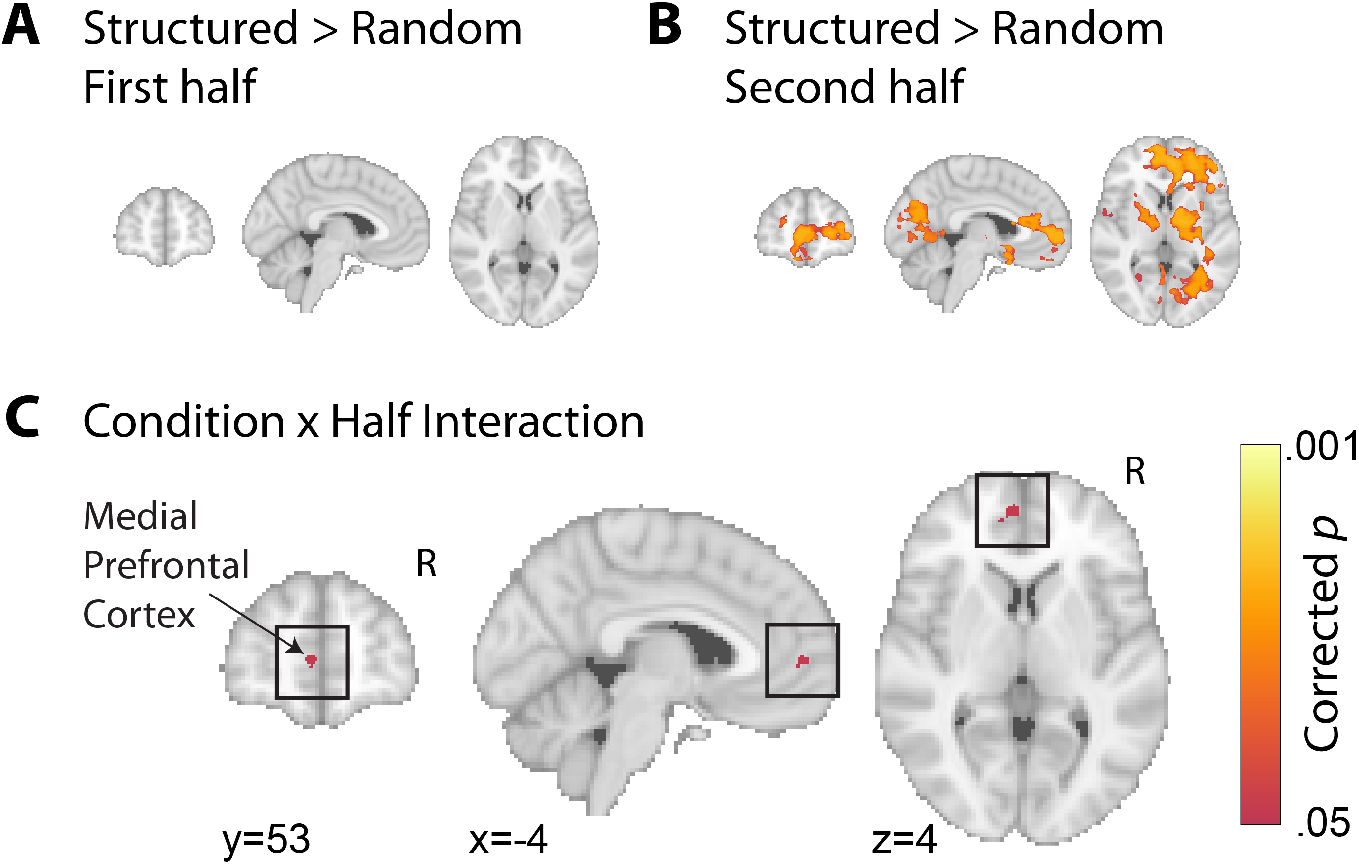
Exploratory whole-brain analysis. Voxelwise contrast of BOLD activity between Structured and Random blocks in (A) the first half and (B) the second half. (C) The medial prefrontal cortex (mPFC) showed an interaction between condition and half, with a greater difference between Structured and Random in the second vs. first half. Voxels in color were significant after correction for multiple comparisons (threshold-free cluster enhancement, one-tailed corrected *p*<.05). Coordinates are in MNI space.

**Table S1.** Demographic information. ‘ID’ is a unique infant identifier (i.e., sXXXX_Y_Z), with the first four digits (XXXX) indicating the family, the fifth digit (Y) the child number within family, and the sixth digit (Z) the session number with that child. ‘Age’ is recorded in months. ‘Sex’ is female or male. ‘Site’ is Scully Center for the Neuroscience of Mind and Behavior at Princeton University (P.ton), Magnetic Resonance Research Center at Yale University (MRRC), or Brain Imaging Center at Yale University (BIC). ‘1st half’ is the number of usable blocks from the first half of exposure (max 6). ‘2nd half’ is the number of usable blocks from the second half of exposure (max 6). ‘Start cond.’ is the randomly selected condition of the first block (alternating thereafter). ‘TR prop’ is the proportion of TRs included from usable blocks. ‘Eye prop’ is the proportion of eye tracking data included from usable blocks. ‘Eye IRR’ is the proportion of frames coded the same way across gaze coders; one participant without eye-tracking data has “nan”.

| ID | Age | Sex | Site | 1st half | 2nd half | Start cond. | TR prop | Eye prop | Eye IRR |
| --- | --- | --- | --- | --- | --- | --- | --- | --- | --- |
| s2307_1_1 | 19.9 | M | P.ton | 4 | 4 | Random | 0.97 | 0.90 | 0.86 |
| s0307_1_2 | 9.1 | M | P.ton | 6 | 2 | Structured | 0.91 | 0.94 | 0.92 |
| s8187_1_8 | 23.1 | F | P.ton | 6 | 2 | Random | 0.92 | 0.83 | 0.95 |
| s2307_1_2 | 21.7 | M | P.ton | 6 | 4 | Random | 1.00 | 0.90 | 0.94 |
| s8187_1_4 | 13.8 | F | P.ton | 4 | 2 | Structured | 0.86 | 0.83 | 0.73 |
| s1187_1_1 | 20.7 | F | P.ton | 6 | 6 | Structured | 0.95 | 0.88 | 0.97 |
| s2687_1_1 | 4.0 | M | MRRC | 3 | 5 | Random | 0.82 | 0.86 | 0.83 |
| s6687_1_1 | 5.0 | F | MRRC | 6 | 4 | Structured | 0.79 | 0.86 | 0.90 |
| s8607_1_1 | 8.5 | F | MRRC | 6 | 2 | Structured | 0.96 | 0.93 | 0.93 |
| s3607_1_1 | 7.5 | F | MRRC | 5 | 7 | Random | 0.88 | 0.91 | 0.96 |
| s8687_1_4 | 14.5 | F | BIC | 6 | 6 | Structured | 1.00 | 0.86 | 0.98 |
| s2687_1_3 | 9.9 | M | BIC | 6 | 2 | Structured | 0.88 | 0.84 | 0.96 |
| s2687_1_4 | 16.3 | M | BIC | 6 | 2 | Structured | 0.86 | 0.89 | 0.90 |
| s6687_1_3 | 11.3 | F | BIC | 6 | 2 | Random | 0.99 | 0.94 | 0.94 |
| s4607_1_4 | 13.0 | F | BIC | 6 | 6 | Structured | 0.95 | 0.88 | 0.93 |
| s4607_1_5 | 14.1 | F | BIC | 6 | 4 | Structured | 0.99 | 0.80 | 0.90 |
| s6607_1_2 | 7.4 | M | BIC | 6 | 6 | Random | 0.97 | nan | nan |
| s0607_1_5 | 17.7 | M | BIC | 6 | 6 | Structured | 0.91 | 0.90 | 0.97 |
| s1607_1_2 | 10.7 | M | BIC | 6 | 6 | Structured | 0.82 | 0.84 | 0.95 |
| s6057_1_1 | 3.6 | M | BIC | 6 | 4 | Random | 0.80 | 0.97 | 0.99 |
| s0057_1_3 | 9.0 | F | BIC | 5 | 3 | Structured | 0.83 | 0.86 | 0.89 |
| s7017_1_1 | 5.2 | F | BIC | 4 | 6 | Random | 0.87 | 0.95 | 0.98 |
| s7017_1_2 | 6.6 | F | BIC | 6 | 6 | Structured | 0.97 | 0.95 | 0.99 |
| s7067_1_1 | 4.7 | F | BIC | 4 | 6 | Structured | 0.96 | 0.95 | 0.95 |
| Av. | 11.55 | . | . | 5.46 | 4.29 | . | 0.91 | 0.89 | 0.93 |

## Notes

### Competing Interest Statement

The authors have declared no competing interest.

